# Detrimental Influence of Arginase-1 in Infiltrating Macrophages on Post-Stroke Functional Recovery and Inflammatory Milieu

**DOI:** 10.1101/2024.07.08.602449

**Authors:** Hyung Soon Kim, Seung Ah Jee, Ariandokht Einisadr, Yeojin Seo, Hyo Gyeong Seo, Byeong Seong Jang, Hee Hwan Park, Won-Suk Chung, Byung Gon Kim

**Affiliations:** Department of Brain Science, Ajou University School of Medicine, Suwon, 16499, Republic of Korea; Neuroscience Graduate Program, Department of Biomedical Sciences, Ajou University Graduate School of Medicine, Suwon, 16499, Republic of Korea; Department of Biological Sciences, Korea Advanced Institute of Science and Technology (KAIST), Daejeon, Republic of Korea; Center for Vascular Biology, Institute for Basic Science (IBS), Daejeon, Republic of Korea; Department of Neurology, Ajou University School of Medicine, Suwon, 16499, Republic of Korea

**Keywords:** Ischemic stroke, Arginase-1, Infiltrating macrophages, Microglia, Synapse elimination, Fibrosis, Functional recovery

## Abstract

Post-stroke inflammation critically influences functional outcomes following ischemic stroke. Arginase-1 (Arg1) is conventionally understood as a marker for anti-inflammatory macrophages, associated with the resolution of inflammation and promotion of tissue repair in various pathological conditions. However, its specific role in post-stroke recovery remains to be elucidated. This study investigates the functional impact of Arg1 expressed in macrophages on post-stroke recovery and inflammatory milieu. We observed a time-dependent increase in Arg1 expression, peaking at 7 days after photothrombotic stroke in mice. Cellular mapping analysis revealed that Arg1 was predominantly expressed in LysM-positive infiltrating macrophages. Using a conditional knockout (cKO) mouse model, we examined the role of Arg1 expressed in infiltrating macrophages. Contrary to its presumed beneficial effects, Arg1 cKO in LysM-positive macrophages significantly improved skilled forelimb motor function recovery after stroke. Mechanistically, Arg1 cKO attenuated fibrotic scar formation, enhanced peri-infarct remyelination, and increased synaptic density while reducing microglial synaptic elimination in the peri-infarct cortex. Gene expression analysis of FACS-sorted microglia revealed decreased TGF-β signaling and pro-inflammatory cytokine activity in peri-infarct microglia from Arg1 cKO animals. In vitro co-culture experiments demonstrated that Arg1 activity in macrophages modulates microglial synaptic phagocytosis, providing evidence for macrophage-microglia interaction. These findings provide new insights into Arg1 function in CNS injury and highlight an interaction between infiltrating macrophages and resident microglia in shaping the post-stroke inflammatory milieu. Our study identifies Arg1 in macrophages as a potential therapeutic target for modulating post-stroke inflammation and improving functional recovery.

## Introduction

CNS injuries following ischemic stroke often result in lifelong functional disabilities (1). While endovascular thrombus removal during the hyperacute stage has significantly improved functional outcomes (2–4), only a small proportion of acute ischemic stroke patients are eligible for mechanical thrombectomy (5, 6). Moreover, a substantial number of treated patients still experience moderate to severe functional impairments (7, 8). Consequently, there is a critical need to develop therapeutic strategies, either as standalone treatments or in combination with endovascular procedures, to mitigate the cellular and molecular effects of ischemic injury during the subacute to chronic stages.

Post-stroke inflammation has emerged as a promising target for modulating functional recovery (9–12). In the acute phase of ischemic stroke, damaged tissues release damage-associated molecular patterns and inflammatory cytokines, activating glial cells such as astrocytes and microglia. This process is accompanied by the infiltration of various immune cells from peripheral blood, including neutrophils, lymphocytes, monocytes, and macrophages, all contributing to the post-stroke inflammatory milieu (13–15). Among these, macrophages and microglial cells, both of myeloid lineage, undergo dynamic phenotypic changes during post-stroke pathological processes. The phenotypic variability of these cells, traditionally categorized as either M1 or M2 at their polarized extremes, plays a crucial role in shaping the inflammatory landscape following stroke (16–18). Recent studies have revealed complex interactions between microglial cells and infiltrating monocytes/macrophages in various CNS injuries (13, 19–21), further complicating our understanding of the inflammatory environment and its regulation.

Arginase-1 (Arg1), a canonical marker for M2 macrophages, is known for its role in resolving pro-inflammatory microenvironment (22). As an enzyme that converts L-arginine to L-ornithine in the urea cycle (23), Arg1 plays a pivotal role in determining macrophage phenotypic characteristics, particularly given that arginine is also a substrate for inducible nitric oxide synthetase, which generates oxidative stress (24, 25). The functional contributions of Arg1 in macrophages have been studied in various inflammatory conditions in peripheral organs outside of the brain. For instance, Arg1 activity is required for suppressing chronic inflammation and promoting wound healing in cutaneous injuries (26). In the kidney, Arg1 is required for macrophage-mediated renal tubule regeneration (27). However, few studies have investigated the functional contribution of Arg1 in macrophages to post-stroke inflammation and functional recovery. The present study revealed that, contrary to the traditional view of Arg1 as a marker for macrophages promoting tissue repair (28), Arg1 in macrophages plays a detrimental role in motor functional recovery following stroke. Our results indicate that Arg1 may induce excessive fibrosis at the border between the infarction core and peri-infarct spared cortical tissue and increase pathological synaptic elimination by altering macrophage-microglial interaction. These findings suggest that Arg1 may be a promising therapeutic target for enhancing functional recovery through modification of the post-stroke inflammatory milieu.

## Materials and methods

### Experimental animals

All animal experiments were performed under the approval of the Institutional Animal Care and Use Committee of Ajou University Medical Center. Arg1 flox (C57BL/6-Arg1tm1Pmu/J, Strain #:008817), Cx3cr1-GFP (B6.129P2(Cg)-Cx3cr1tm1Litt/J, Strain #:005582), Rosa-stop-eYFP (C57BL/6Gt(ROSA)26Sortm1(HBEGF)Awai/J, Strain #:007900), and LysM-cre (B6.129P2-Lyz2tm1(cre)Ifo/J, Strain #:004781) lines were purchased from Jackson laboratory. Animals were maintained in individually ventilated cages with 12 hours of light/dark cycle. To create a conditional knockout (cKO) of Arg1 animals, specifically in cells expressing LysM, Arg1 flox mice were bred with LysM-cre mice, where the Cre recombinase is expressed under the control of LysM promoter. Animals that were homozygous for the Arg1 flox allele and heterozygous for the LysM-cre allele were designated as Arg1 cKO mice. Arg1 flox homozygous animals were used as controls. To generate mice expressing YFP under the LysM promoter control, we crossed Rosa-stop-eYFP mice with LysM-cre line. Mice heterozygous for both alleles were used in experiments.

### Induction of Photothrombotic ischemic stroke

Animals were anesthetized by intraperitoneal injection of ketamine (90mg/kg) and xylazine (10mg/kg) mixture before surgery. The head was secured in the stereotaxic frame, and an incision was made to expose the skull after a brief sterilization using 70 % ethanol. A cold light source, utilizing a 150W halogen lamp fiberoptics with a diaphragm of 1.5 mm diameter, was placed on either side of the skull corresponding to the caudal forelimb area at the predetermined coordinates (AP 0.0 mm, ML ± 1.8 mm). For animals subjected to behavioral assessment, illumination was directed to the side contralateral to the preferred forelimb determined during the shaping session for the pellet retrieval test. Animals not subjected to the behavioral study underwent photothrombosis induction in the right hemisphere. Rose Bengal (Sigma-Aldrich, 15 μg/ml) was administered intraperitoneally, and illumination started 5 minutes after injection. The cold light was illuminated continuously for exactly 25 minutes. To prevent hypothermia, animals were maintained on a 37°C heating pad throughout surgery and illumination. Animals in the sham group received an intraperitoneal injection of PBS vehicle instead of Rose Bengal, along with the same duration of light illumination.

### Behavioral assessment of motor functions

#### Pellet retrieval test

The pellet retrieval test was performed using a custom-made apparatus applying previously reported protocol (29). The apparatus, measuring 15 cm wide, 8.5 cm long, and 20 cm high, was created for single or double slits so that mice can retrieve food pellets with their forelimbs. In the single-slit configuration, the opening was centrally positioned, allowing mice to use either forelimb. The double-slit setup placed openings near the side walls, forcing mice to use only their dominant forelimb to reach the food pellets. To motivate animals to consume the sucrose precision pellets (20 mg, Bio-Serv), during the pellet retrieval test, they were fasted by providing only 10% of their body weight in regular chow daily. During the initial two days of fasting, precision pellets equivalent to 10% of the chow weight were provided to help the animals recognize the precision pellet as an energy source. This regimen allowed the animals to maintain approximately 80% of their body weight during the behavior test. Following a 2-day fasting, animals were habituated to the reaching task apparatus for another 2 days. On the first day of habituation, two animals were habituated simultaneously for 20 minutes each with pellets. On the second day, single animals were habituated for 20 minutes each. During this shaping session, the single-slit configuration was used so that the animals could freely reach for them using their forelimbs through a 0.5 mm single slit. Only animals that consumed over 20 pellets within 20 minutes and exhibited over 70% hand preference were subjected to the subsequent pre-training. Pre-training consisted of 7 consecutive days during which the animals consumed 40 pellets per day using the double-slit configuration. The baseline for the reaching task was established on the last day of pre-training. Behavioral evaluation was conducted every 7 days following the ischemic stroke.

Animals underwent another 2-day fasting period before each behavior session. The success rate of the pellet retrieval task was measured by dividing the number of successful pellet retrievals by the total number of pellets consumed. A successful pellet retrieval was defined as the animal directly consuming the pellet after grasping it. If the pellet was displaced and could not be reached by the forelimb or if the animal dropped it before consuming it, the trial was considered unsuccessful.

#### Ladder rung test

Pre-training for the ladder rung test was initiated on the third day of pre-training for the pellet retrieval test and continued over 5 consecutive days. Animals traversed a custom-made ladder, measuring 60 cm in length with 20 rungs spaced 1.0 cm apart, for five crossings per day. One precision sucrose pellet was positioned on each side of the ladder for every crossing attempt to motivate the animals. The foot fault rate was determined by counting the number of misplaced forelimbs and dividing it by the total number of steps taken on each side of the ladder. The foot fault rate was averaged across the five ladder crossings for each animal.

#### Cylinder test

The cylinder test was performed using a custom-made plexiglass cylinder which was 17.5 cm height and 8.8 cm inner diameter. Perpendicular mirror was placed at the background to record forelimb touches against the camera. The baseline for forelimb usage in the cylinder test was assessed on the final day of behavioral training without prior acclimatization. The forelimb usage ratio was calculated after observing at least 40 forelimb touches against the cylinder wall. The number of touches made by both impaired and non-impaired forelimbs was recorded. The forelimb usage ratio was calculated by dividing the number of touches made by the impaired limb by the number made by the non-impaired limb.

### mNSS scoring

The baseline for the modified neurological severity score (mNSS) was evaluated one day before inducing the stroke. mNSS was determined as previously described (30). A customized apparatus was constructed for the beam test of mNSS scoring, where animals were allowed to walk on a 1.0 cm-width plexiglass beam.

### Tissue preparation and immuno-histochemistry

Animals were euthanized via transcardiac perfusion using cold phosphate-buffered saline (PBS) followed by 4% paraformaldehyde (PFA). Following perfusion, brain tissue was extracted for histological analysis. The tissue was post-fixed in 4% PFA overnight. The next day, the tissue was immersed in a 30% sucrose solution until it sank to the bottom for cryoprotection. Subsequently, the tissue was sectioned to 30 μm thickness using a Cryo-Microtome. The sections were then placed in a glycine solution composed of 30% glycerol, 30% ethylene glycol, 30% distilled water, 10% PB buffer. The tissue sections were then transferred onto glass slides for further immunolabeling procedures. The following primary antibodies were applied after dilution in a 10% NGS solution overnight at 4°C, following 30 minutes of tissue penetration in a 10% NGS, 0.3% Triton X-100 solution at room temperature: anti-Arg1 c-terminal (Abcam, 60176, 1:500), anti-Iba-1 (Wako, 019-19741, 1:500), anti-GFAP (DAKO, z0034, 1:1000), anti-fibronectin (Merck, MAB1940, 1:500), anti-CSPG (CS56, Millipore, C8035), anti-vGlut2 (Synaptic Systems, 135416, 1:500), anti-PSD95 (Invitrogen, 1:200), anti-MAP2 (Millipore, ab5622 1:500), anti-MBP (Millipore, AB980, 1:1000), anti-Ly6G (Invitrogen, 14-5931-81, 1:200), and anti-GFP (Abcam, 13970, 1:500). For the labeling of perineuronal nets, biotinylated Wisteria floribunda agglutinin (Vector lab, b-1355) was used. Secondary antibodies were applied for 2 hours at room temperature. Tissues were briefly washed with PBS, and coverslips were mounted on the slides using DAPI-containing mounting solutions.

### Image acquisition and analysis

Tissue images were acquired using LSM800 (Zeiss, German) confocal microcopy. For imaging of synapses and Iba-1 3D constructions, Nikon A1R (Nikon, Japan) confocal microscopy was used. Images were captured using a 100x oil immersion objective lens. Following image acquisition, the images were analyzed by ImageJ software (https://imagej.net/ij/index.). All regions of interest (ROI) were subjected to channel separation processing to quantify the intensity of the images. Images were background subtracted and converted into binary images. The intensity of duplicated original images was measured based on the mean grey values derived from masked binary images. All images were processed with the same parameters. The average mean grey value was utilized to measure immunolabeling fluorescent intensity.

### Tissue protein extraction and western blot analysis

Protein samples were extracted from the tissue after homogenization in tissue lysis buffer (T-per supplemented with phosphatase/protease inhibitor). Samples were centrifuged at 15000 g for 5 minutes at 4 °C. Electrophoresis was performed at 60 V for 30 minutes followed by 100 V for 60 minutes. After blocking with 5 % skim milk, primary antibodies against Arg1 (Abcam, Ab91279, 1:1000) diluted in 5 % skim milk were incubated overnight at 4 °C with horizontal agitation. After washing, the membranes were incubated with a secondary antibody for 2 hours at room temperature. For loading control, HRP-conjugated anti-beta-actin (Sigma-Aldrich, A3850, 1:20000) was incubated for 1 hour at room temperature. The membranes were then developed and imaged using a Gel doc system (Bio-rad) after applying HRP substrates for 1 minute at room temperature.

### *In vivo* phagocytosis analysis and 3D image reconstruction

To analyze in vivo synaptic elimination by microglia after stroke, AAV9-hSYN-mCherry-eGFP was injected into the rostral forelimb area of the premotor cortex (AP −1.5 mm, ML −1.8 mm, DV −1.5 mm) of mice. Two weeks after viral injection, photothrombotic stroke was induced in the caudal forelimb area. The brain was sectioned into 30 μm-thickness and labeled with Iba-1 antibody to visualize microglia. At least 5 regions of interest (ROIs) were imaged approximately 360 μm away from the stroke boundary using a Nikon A1R confocal microscope. The image files were converted to IMARIS 9.0 software formats for 3D reconstructions. The phagocytosed synapsed Iba-1 signals were reconstructed into 3D surfaces, while mCherry or eGFP signals were converted into 3D spots. Rendered 3D spots outside the Iba-1 3D surface were removed. To detect mCherry signals alone within 3d rendered Iba-1, any mCherry signal co-localized with eGFP was masked. Although mCherry and eGFP+ colocalized signals were also detected in Iba-1 cells, the majority of signals were from mCherry alone. eGFP alone spots were rarely detected in Iba-1 positive cells. Non-colocalizing mCherry spots with Iba-1 surface were considered as “phagocytosed synapses”. The number of phagocytosed synapses was analyzed from 136 or 137 Iba-1+ cells from Arg1 flox or Arg1 cKO groups, respectively. Individual cell values and averaged group values were plotted separately.

### Single cell dissociation and Fluorescent activated cell sorting (FACS) cell sorting

To isolate microglia from the peri-infarct cortex, freshly isolated brain was placed on a brain matrix, and the brain tissue encompassing the infarcted and spared peri-infarct cortex was dissected into pieces measuring 3.0 x 3.0 mm. Then, the infarct core was carefully dissected and removed using a razor blade. Subcortical structures, including the white matter, were removed using a razor blade. For sham animals, tissue from the same location and size was collected. After tissue collection, single cell dissociation was performed using the Adult brain dissociation kit (Miltenyi Biotec, #130-107-677) according to the manufacturer’s instructions of the single cell dissociation kit. Briefly, tissues were enzymatically digested in the gentle MACS dissociator (Miltenyi Biotec, #130-093-235) for 30 minutes at 37 °C. Following dissociation, the tissue suspensions were pelleted and resuspended in debris removal solutions. Dulbecco’s phosphate-buffered saline (D-PBS) was layered over the tissue suspension in debris removal solutions and centrifuged at 4°C for 10 minutes. After centrifugation, tissue debris and myelin debris were removed from the tissue gradient. The resulting tissue pellets were then resuspended in red blood cell (RBC) removal solutions and centrifuged once more. After discarding the supernatants, the tissue pellets were resuspended in fluorescence-activated cell sorting (FACS) buffer, consisting of PBS supplemented with 1% bovine serum albumin (BSA) and 0.1% sodium azide. Cell numbers were determined using a LUNA cell counter (Logos bios, South Korea). For FACS antibody labeling, anti-CD11b-Alexa488 (Bio-rad, BRMCA275A488) and anti-CD45-Alexa647 (Bio-rad, BRMCA43A647) antibodies were incubated (10 μl per 1.0 x 10^6^ cells) for 30 minutes in an ice bucket covered with aluminum foil to protect the samples from light exposure. Following incubation, cells were centrifuged and resuspended in PBS to wash off unbound antibodies. Cell sorting was performed using a FACS Aria III (BD), with gating strategies based on control IgG fractions. based on control IgG fractions. To isolate microglial fractions, CD11b-positive and CD45-low populations were collected into 1.5 ml Eppendorf tubes. To preserve RNA integrity for subsequent RNA extraction procedures, the sorted samples were immediately immersed in Qiazol solution.

### Cytokine PCR array and bioinformatics analysis

Total RNA was extracted using Qiazol (Qiagen) according to the manufacturer’s instructions. RNA concentration was determined using a Nanodrop spectrophotometer (Thermo Scientific). cDNA synthesis was performed using the RT² First Strand Kit (Qiagen). The RT² Profiler™ PCR Array Mouse Common Cytokine (Qiagen, PAMM-021ZA-12) was used to analyze cytokine gene expression, following the manufacturer’s protocol. The samples were loaded into the 96-well format of the array plate. Amplification was conducted using the QuantStudio 3 Real-Time PCR System (Thermo Fisher Scientific). Raw data were extracted from the QuantStudio 3 software and analyzed for gene expression changes. To identify hub cytokine genes that were significantly affected in microglial cells within the peri-infarct tissue, we performed STRING analysis (https://string-db.org/). A list of genes with their corresponding fold changes, up-regulated and down-regulated, at a significance level of *p* < 0.05 was generated on the webpage to visualize interactions among the listed genes. The parameters used for STRING analysis were specified as follows: full STRING network type, interaction sources included text mining, experiments, and databases, and the minimum required interaction score was set to 0.400 (medium). The maximum number of interactors displayed was limited to the 1st shell with 5 interactors. The web software performed K-means clustering on the results, which were then used to calculate protein interactions by dividing the number of nodes of each gene by the total number of listed genes (node degree/total nodes).

### Real-time RT-PCR

One microgram of total RNA was used for cDNA synthesis using CellScript cDNA synthesis kit (CellSafe, South Korea). After preparing cDNA mixture, samples were incubated for 5 minutes at 95 °C followed by 60 minutes at 60 °C using Thermocycler (Bio-rad, U.S). The samples were stored in −20 °C until use. RT-PCR was performed using Accupower PCR Premix (Bioneer, K2016) supplemented with 1.0 μl of cDNA and primers. One μl of cDNA samples was mixed with SYBR, ROX dye, and the following primers. 18S: Forward: 5’- CGCGGTTCTATTTTGTTGGT-3’, Reverse: 5’-AGTCGGCATCGTTTATGGTC-3‘, Arg1: Forward: 5’-AGGAGCTGTCATTAGGGACATC-3’, Reverse: 5’-CTCCAAGCCAAAGTCCTTAGAG–3, trem2: Forward: 5’-CTTGCTGGAACCGTCACCAT-3’, Reverse: 5’-CTCTTGATTCCTGGAGGTGC-3‘, gal-3: Forward: 5’-CTACCCAGGGGCTGCTTATC-3’, Reverse: 5’-AGCGGGGGTTAAAGTGGAAG-3‘, cd36: Forward: 5’-TGGAGGCATTCTCATGCCAG-3’, Reverse: 5’-AAGACACAGTGTGGTCCTCG-3, cd68: Forward: 5’-GGGCTCTTGGGAACTACACG-3’, Reverse: 5’-AGACTGTACTCGGGCTCTGA-3‘, cr3: Forward: 5’-AAGCAGCTGAATGGGAGGAC-3’, Reverse: 5’-TAGATGCGATGGTGTCGAGC-3‘. Amplifications were performed using QuantStudio3 PCR amplifier. The Ct values of amplified genes were analyzed using QuantStudio software. Raw data were calculated as relative mRNA fold changes by comparing the Ct value of the endogenous 18S housekeeping gene.

### Cell culture

To obtain bone marrow-derived macrophages (BMDMs), a C57BL6/N mouse was sacrificed in a CO2 chamber. The tibia and femur were dissected from both hindlimbs, and the bone marrow was flushed with ice-cold PBS and collected in DMEM on ice. The bone marrow was centrifuged at 300 g for 5 minutes and resuspended in DMEM containing 10% FBS and 1% penicillin/streptomycin. The cell suspension was filtered through a 70 μm cell strainer and seeded into a 150 mm petri dish with 40 ng/ml of m-CSF (BioLegend, 576406) for 7 days. For microglial culture, mixed glial cells were obtained from the brains of C57BL6/N P1 pups. Briefly, P1 mice were sacrificed by decapitation, and their whole brains were collected in ice-cold DMEM and mechanically homogenized. The homogenate was filtered through a 70 μm cell strainer. After centrifugation at 300 g for 5 minutes, the pellet was resuspended in DMEM containing 10% FBS and 1% penicillin/streptomycin. Two P1 pup brains were used to culture mixed glia in a 75 mm culture flask. To collect microglia from the mixed glial culture, the flask was incubated on a horizontal shaker at 200 rpm for 90 minutes 14 days after the initial culture. The culture medium was collected after shaking and centrifuged at 300 g for 5 minutes to collect the microglia. The cell pellet was resuspended in serum-free DMEM, and 5 x 10^5^ microglia were seeded on a PDL pre-coated 6-well plate.

To establish microglia-BMDM co-culture, BMDMs were enzymatically detached using 0.25% trypsin (Gibco, SH30042.01) 7 days after initial seeding. 5 x 10^5^ cells were seeded on a 6-well culture insert (Falcon, 353090, 0.4 μm pore). After treatment with pharmacological agents, Zymosan (Sigma, Z4250, 1.25 μg/ml) and/or OATD-02 (Molecure, 1.0 μM), the BMDM culture insert was transferred to a 6-well plate containing primary microglia and co-cultured for 24 hours. Then, microglial cells were either harvested for mRNA extraction or subjected to *in vitro* phagocytosis assay.

### *In vitro* microglia synaptosome phagocytosis assay

To isolate synaptosome preparations, cortical tissue (~10.0 mg) was dissected from the brain of C57BL6/N wild-type mice. The tissue was immersed in 100 μl of 0.32M sucrose and homogenized using a tissue homogenizer. The homogenate was centrifuged at 470 g for 2 minutes, and the supernatant was collected. The supernatant was then centrifuged again at 10,000 g for 10 minutes. The resulting supernatant was removed, and the pellet was resuspended in 0.32M sucrose solution. This suspension was transferred into 0.8M sucrose solution and centrifuged at 9100 g for 15 minutes using a swing bucket rotor. After centrifugation, the synaptosome layer between the 0.32M and 0.8M sucrose solutions was collected. To label the synaptosomes with pHrodo, they were resuspended in sodium bicarbonate buffer (pH 8.4) at a concentration of 3 mg/ml. pHrodo (Invitrogen, P36600, 3 μM) dissolved in DMSO was added to the synaptosomes and incubated for 45 minutes at room temperature. The synaptosomes were then diluted in DPBS (Gibco, 14190-144) at a 1:10 ratio and centrifuged for 5 minutes at 2500 g, twice, to wash away excess pHrodo. The synaptosomes were resuspended in serum-free DMEM at 3 mg/ml concentration and used for the in vitro phagocytosis assay. After 24 hours of co-culture with BMDMs, pHrodo-labeled synaptosomes were added to the microglia at a final 0.017 mg/ml concentration. The cell culture plate was placed in the JULI stage (NanoEnTek) and live images were taken for 24 hours at 80-minute intervals. Five regions of interest (ROIs) from each well were imaged using a 10x lens. To analyze the number of pHrodo-positive microglia, the JULI STAT software (ver. 2.0.1) was used, which automatically counted the number of pHrodo-positive cells at each time point.

### Arginase activity assay

Enzyme activity was measured using Arginase Enzyme Activity Assay Kit (Sigma-Aldrich, MAK112) To measure Arg1 activity in cultured BMDMs, 150 μl of 0.01 M tris-HCl (pH 7.4) was added. After incubation for 10 minutes at room temperature, the cell lysate was centrifuged for 10 minutes at 500 g to remove cell debris. Urea standard (50 mg/dL), lysis buffer, water, cell lysates were added to a 96-well plate. following the addition of arginine substrate buffer (20 % Mn solution), the plate was incubated for 2 hours at 37 °C. After incubating for 2 hours, UREA reagent was added to each well and incubated for 1 hour at room temperature. the absorbance at 430nm was measured to calculate the arginase activity of the cell lysate.

### Statistical analysis

Statistical analyses were conducted using GraphPad Prism software (Ver.9.0). An unpaired t-test was performed to compare two independent groups. One-way ANOVA was used to compare more than two independent group. Behavioral assessments were analyzed using Two-way ANOVA to account for the time difference within the same subject. Turkey’s post-hoc analysis was performed for the statistical hypothesis test.

## Results

### Arg1 expression following photothrombotic stroke in mice

Arg1 protein expression was assessed by immunohistochemistry at different time points following the induction of a photothrombotic infarction in mice. The photothrombosis protocol used in this study generated a localized infarction, affecting the entire thickness of the cerebral cortex while preserving the corpus callosum (Figure 1A). Arg1-positive cells were observed within the infarcted tissue in a time-dependent manner. Arg1 expression was first noticeable 3 days post-stroke (dps) and peaked at 7 dps (Fig. 1B, C). By 14 dps, the number of Arg1-positive cells decreased almost to the baseline level (Figure 1B, C). The Arg1-expressing cells were mainly distributed at the periphery of the infarction core at 7 dps when the expression level was at its highest (Fig. 1D). Arg1 expression was not observed in peri-infarct neurons visualized by microtubule-associated protein 2 (MAP2) (Fig. 1E). The expression of Arg1 was restricted within the GFAP-positive astrocytic scar boundary (Fig. 1E). The majority of Arg1-expressing cells were co-localized with Iba-1-positive macrophages or microglia (Fig. 1E), indicating that Iba-1-positive myeloid cells were the main source of Arg1 expression after an ischemic stroke. When a photothrombotic infarction was induced in the CX3CR1-GFP mice where GFP is expressed in CX3CR1 positive resident microglial cells (31), only 20% of Arg1 positive cells were colocalized with GFP (Fig. 1F, H). Lysozyme M is expressed exclusively in hematogenous macrophages and neutrophils but not in microglia (32). To determine whether macrophages infiltrating from blood express Arg1 (32), we employed an R26-stop-EYFP reporter line crossed with LysM-cre mice. We observed that approximately 90% of Arg1-positive cells colocalized with YFP-positive cells (Fig. 1G, H). Since LsyM is also expressed in almost half of the neutrophils (32, 33), we also investigated whether Ly6G-positive neutrophils express Arg1. Ly6G-positive neutrophils were observed within the infarcted tissue at 1 and 3 dps when the neutrophils actively infiltrate the brain parenchyma (34). However, Arg1 was not expressed in the Ly6G-positive neutrophils at either time point (Supplementary figure 1. A-C). Collectively, immunolabeling studies suggest that LysM+ macrophages are a principal source of Arg1 after ischemic stroke.

**Figure 1.**
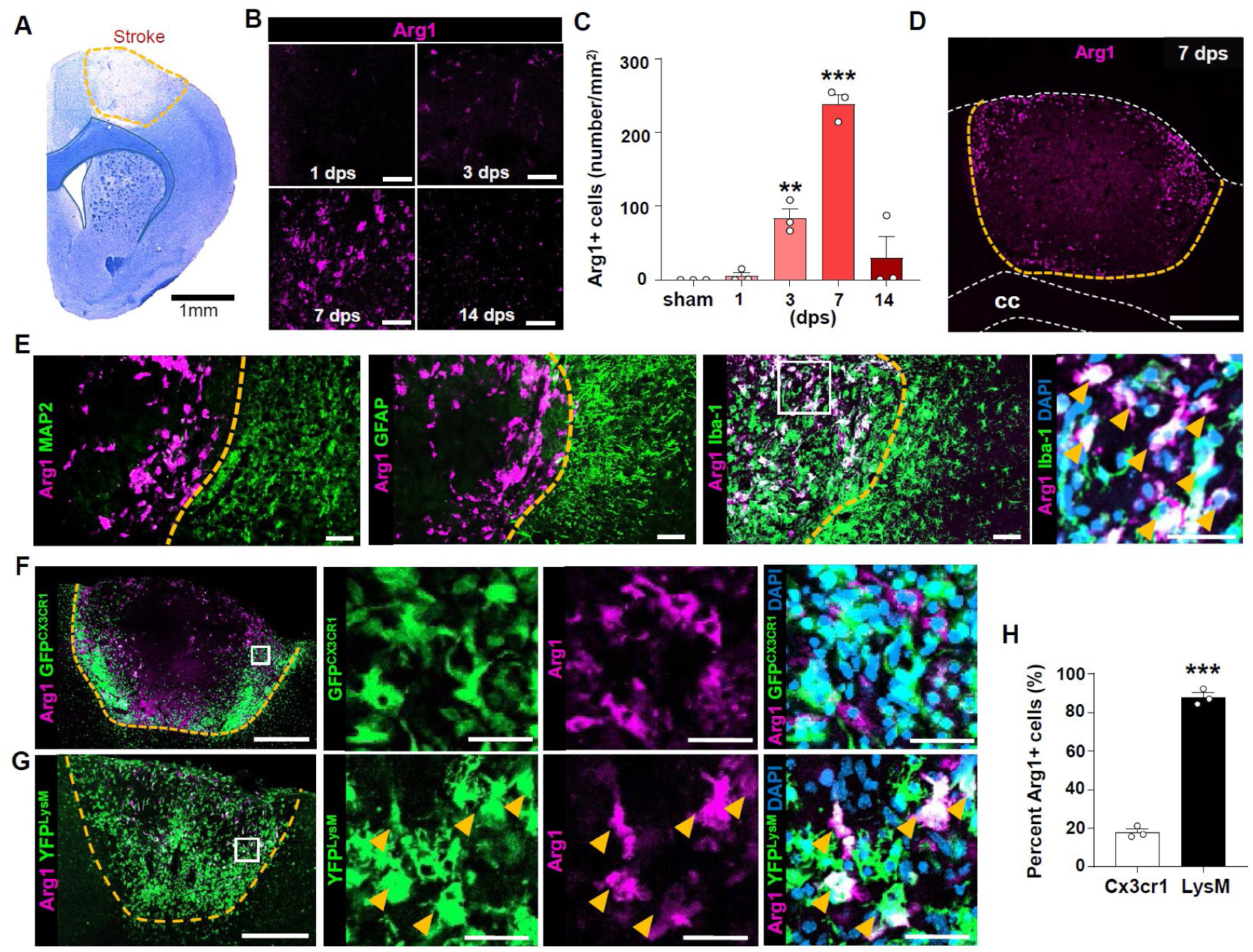
Characterization of Arg1 expression after photothrombotic stroke in mice. (A) Eriochrome cyanine R staining of brain tissue at 7 days post-stroke (dps), showing cortical ischemic region. Scale bar = 1.0 mm. (B) Immunohistochemistry of Arginase-1 (Arg1) at the lesion core at 1, 3, 7, and 14 dps. Scale bar = 50 μm. (C) Quantification of the number of Arg1 positive cells. *, **, and *** indicate *p* < 0.05, *p* < 0.01, and *p* < 0.001, respectively, by one-way ANOVA followed by *post hoc* Bonferroni’s multiple comparison test. N = 3 / time points. (D) A representative image show localization of Arg1 positive signals in the periphery of lesion core at 7 dps. Lesion border is indicated in yellow dotted line. Corpus callosum and cortex is indicated in white dotted lines. Scale bar = 500 μm. cc: corpus callosum. (E) Representative immunohistochemistry images of Arg1 and MAP2, GFAP, Iba-1, respectively. Yellow dotted line indicates lesion border. The right-most magnified images were obtained from the white solid squares in Iba-1 and Arg1 co-labeled image. Scale bar = 50μm. (F, G) Immunohistochemistry of Arg1 in Cx3cr1-GFP (F) or LysM-cre::R26-stop-YFP (G) animals. Scale bar = 500 μm for low magnification images and 50 μm for high magnification images. White solid squares indicate regions that are magnified at the right side. (H) Quantification of the percent Arg1 expressing Cx3cr1+ or LysM+ cells out of the total Arg1+ cells at 7 dps. *, **, and *** indicate *p* < 0.05, *p* < 0.01, and *p* < 0.001, respectively, by unpaired T-test.

### Deletion of Arg1 in macrophages promotes recovery of motor function following photothrombotic stroke

To investigate the functional role of Arg1, we generated Arg1 conditional knock-out (cKO) mice by crossing Arg1^flox^/^flox^ and LysM-cre mice lines, aiming to selectively delete Arg1 in LysM-expressing macrophages. In this model, the Cre recombinase, expressed under the control of the LysM promoter, targets the two loxP sequences inserted upstream of exon 7 and downstream of exon 8, respectively, of the Arg1 genomic sequence. It then removes the genomic sequence between the two loxP sites, resulting in the deletion of exons 7 and 8 (Supplementary fig. 2A), which encompass the substrate binding region of Arg1 (amino acids 126-277). Induction of Arg1 mRNA in the brain tissue after stroke was almost completely abolished at 7 dps in Arg1 cKO mice (Supplementary fig. 2B, C). We confirmed that Arg1 protein levels were substantially reduced following stroke in Arg1 cKO mice (Fig. 2A). The number of Arg1 positive cells was markedly decreased in Arg1 cKO mice, although a small proportion of Arg1-expressing cells were still detected in immunohistochemistry within the infarcted region (Fig. 2B, C).

**Figure 2.**
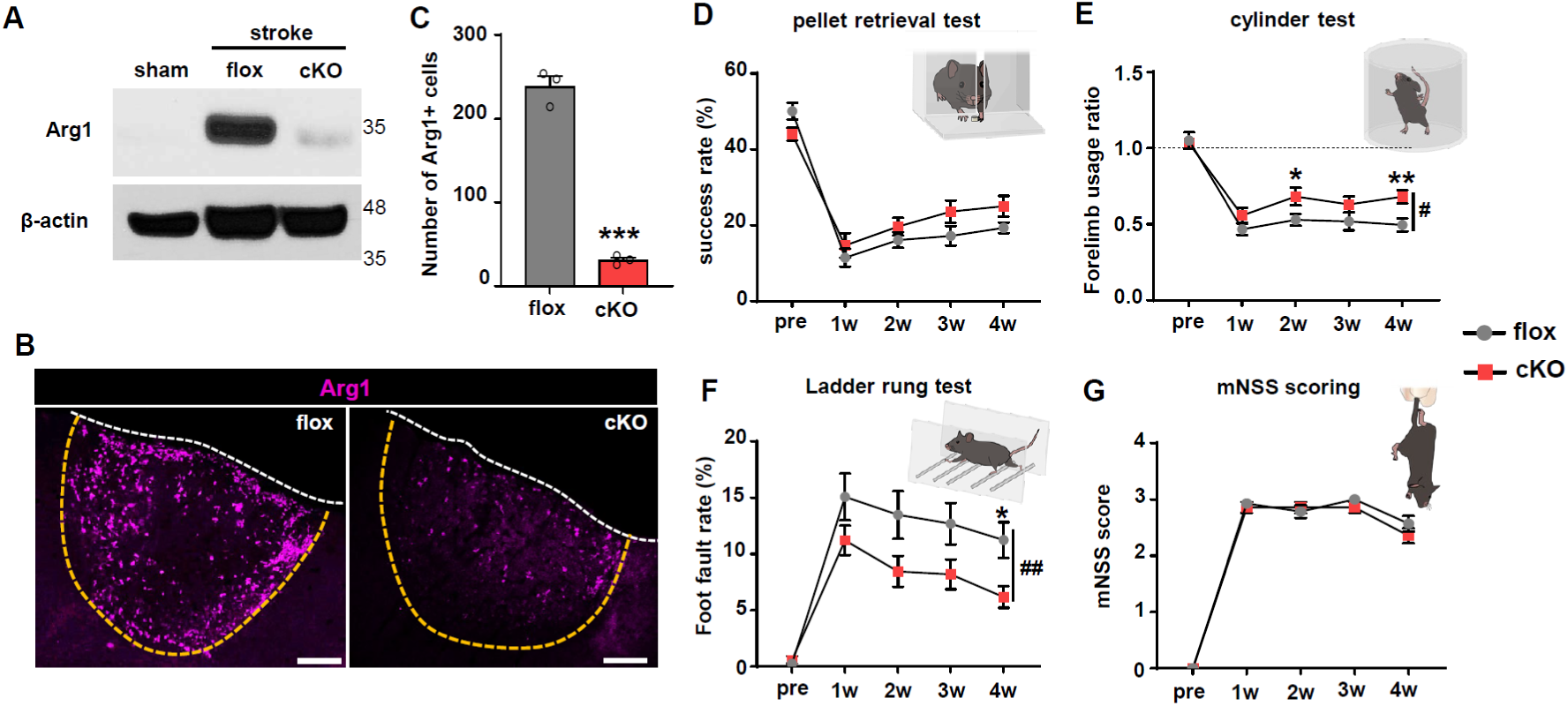
Motor behavior recovery in Arg1 cKO animals after photothrombotic stroke. (A) Western blot of Arg1 protein in the brain tissue at 7 days post-stroke (dps). (B) Immunohistochemistry of Arg1 in Arg1 cKO animals or flox (Arg1 flox/flox) control animals at 7 dps. Scale bar = 200 μm. Infarction: yellow dotted line, cortex: white dotted line. (C) Quantitative analysis of the number of Arg1-expressing cells per mm2. *** indicates p < 0.001 followed by Unpaired T-test. (N = 3 per group). (D-G) Quantitative analysis of pellet test, cylinder test, ladder walk test, mNSS scoring, respectively. Two-way ANOVA repeated measure was performed to match the time difference. Bonferroni’s multiple comparison was conducted to assess group differences at each time point. (N = 14 per group). * indicates p < 0.05 by two-way ANOVA followed by *post hoc* Bonferroni’s multiple comparisons. #, ## indicates p < 0.05, p < 0.01, respectively, by two-way ANOVA column factor (overall group difference).

The animals were subjected to a battery of behavior tests to assess the recovery of motor functions. In the pellet retrieval test, Arg1 cKO animals tended to achieve a higher rate of successful pellet retrieval than control Arg1 floxed animals, although the genotype effect on the successful retrieval rate was not statistically significant according to the two-way repeated measures ANOVA (*F*_(1, 26)_ = 1.309, *p* = 0.263) (Fig. 2D). In the cylinder test, mice without stroke uses both forelimbs equally to touch the cylinder wall. After stroke, they started to preferentially use non-impaired forelimb, and the ratio between impaired and non-impaired limbs became approximately 0.5 at 7 dps. Arg1 cKO mice began to exhibit more frequent use of the impaired forelimbs starting from the 2 week-time point, resulting in a higher ratio than Arg1 flox animals (Fig. 2E). Two-way repeated measures ANOVA revealed a significant difference between the control and cKO group over time (*F*_(1, 26)_ = 5.520, *p* = 0.028). In the ladder rung test, control Arg1 floxed animals made frequent mistakes at 7 dps, and the number of errors declined thereafter. Arg1 cKO animals exhibited much faster recovery in this test, showing lower percentage errors from the 2-week time point (Fig. 2F), and the genotype effect on the difference in the error rate was also statistically significant (*F*_(1, 26)_ = 10.35, *p* = 0.004). Semi-quantitative assessment of neurological functions using mNSS (modified neurological severity score) did not reveal a difference between the two groups (Fig. 2G). When the motor functions were assessed in male and female animals separately, Arg1 cKO mice exhibited a similar pattern of improvement in recovery regardless of the sex difference (Supplementary fig. 3), indicating that sex did not influence functional recovery by Arg1 deletion.

### Arg1 cKO reduces post-stroke fibrosis and promotes peri-infarct remyelination

To determine the pathological outcomes that may underlie the enhanced functional recovery in Arg1 cKO mice, we first examined whether Arg1 deletion altered the activation of macrophage and microglial cells. The intensity of IBA-1 immunoreactivities representing both macrophages and microglial cells was comparable within the stroke lesion core between flox and cKO mice (Supplementary fig. 4A, B). Furthermore, IBA-1 positive cells in the peri-infarct cortex, which highly likely represent microglial cells, exhibited similar immunoreactivities (Supplementary fig. 4A, C). We also compared the astroglial reactivity surrounding the infarct core (Supplementary fig. 4D, E). The intensity of GFAP immunoreactivities was not significantly different between flox and Arg1 cKO mice. Moreover, the volume of infarcted tissue delineated by the GFAP-positive astroglial border was also not affected by Arg1 deletion in macrophages (Supplementary fig. 4D, F).

Arginase activity in macrophages is linked to increased fibrosis in peripheral organs (35, 36). Recent studies revealed that ischemic stroke induces fibrotic scarring, which may exacerbate functional recovery and inhibit the formation of post-stroke axonal connections (37, 38). To assess the extent of post-stroke fibrosis, cortical tissues were stained using antibodies against fibronectin, an extracellular matrix protein comprising fibrotic scars. We observed significantly reduced fibronectin expression in Arg1 cKO animals compared to Arg1 flox control animals following stroke (Figure 3 A, B). Chondroitin sulfate proteoglycans (CSPGs) are produced by astrocytic glia and other components of fibrotic scars such as fibroblasts and macrophages (39, 40). CSPGs deposited in injured CNS can contribute to the formation of fibrotic microenvironment (41, 42). We found that the photothrombotic stroke led to a marked accumulation of CSPGs (measured by CS-56 immunoreactivity) at the border of the infarcted tissue as well as the lesion core (Fig. 3C). The CSPG deposition was sharply attenuated in Arg1 cKO animals (Fig. 3C, D), indicating that Arg1 in macrophages may be implicated in the development of post-stroke fibrotic microenvironment.

**Figure 3.**
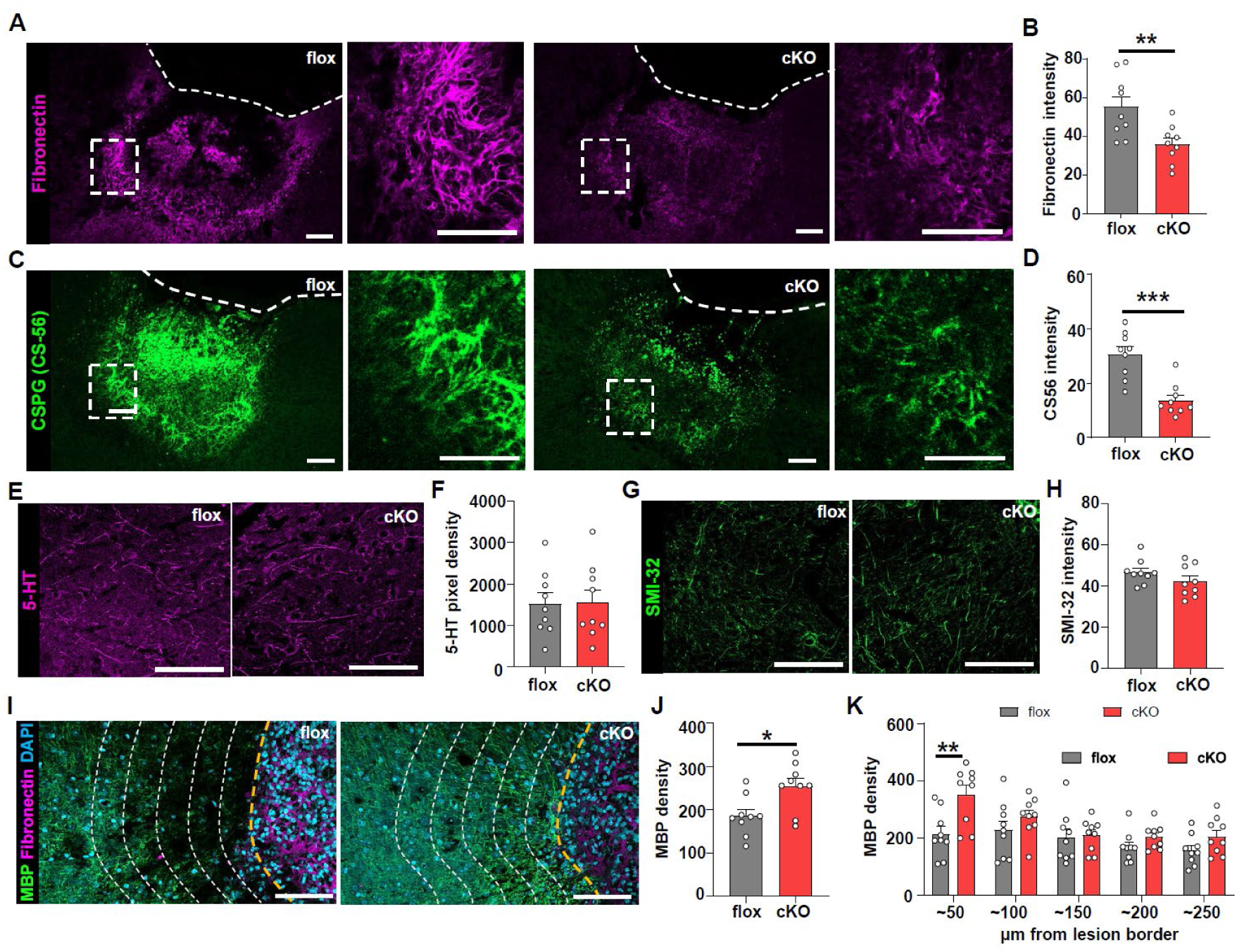
Arg1 cKO reduces post-stroke fibrosis and promotes peri-infarct myelination. (A, C) Representative images of the coronal brain sections subjected to immunohistochemical staining of fibronectin (A) and CS-56 (CSPGs) (C). The tissue sections were obtained from Arg1 cKO or flox control animals sacrificed 4 weeks after a photothrombotic stroke. Dashed rectangles indicate regions that are magnified on the right side. Dashed lines indicate the outer edges of the cerebral cortex. Scale bars = 100 μm. (B, D) Quantification graphs comparing the intensity of fibronectin (B) and CS-56 (D) immunoreactivity. ** and *** indicate *p* < 0.01 and *p* < 0.001, respectively, by unpaired T-test. (E, G) Representative images of the coronal brain sections subjected to immunohistochemical staining of 5-HT (serotonin) (E) and SMI-32 (non-phosphorylated neurofilament) (G). The images were acquired in the peri-infarct cortex. Scale bars = 100 μm. (F, H) Quantification graphs comparing the density of 5-HT (F) and SMI-32 (H) immunoreactivity. (I) Double immunofluorescence staining of myelin basic protein (MBP) and fibronectin. Scale bars = 100 μm. Yellow dashed lines indicate the infarction borders. White dashed lines spaced at 50 μm intervals indicate boundaries for quantification based on distance from the border in (K). (J) A quantification graph of MBP immunoreactivity at the peri-infarct cortex. * indicates *p* < 0.05 by unpaired T-test. (K) A quantification graph of MBP immunoreactivity based on distance from the infarct border. ** indicates *p* < 0.01 by unpaired T-test.

Fibrotic scar acts as a strong barrier that impedes axonal growth following CNS injury (38, 43, 44). When we visualized the serotonergic 5-HT axons in the peri-infarct cortex, the density of the 5-HT axons in the peri-infarct regions was not affected by Arg1 cKO in macrophages (Fig. 3E, F). We also examined the non-phosphorylated neurofilament immunoreactivity (SMI-32) which represents damaged axons (45). The SMI-32 immunoreactivity did not change significantly in the peri-infarct area by Arg1 deletion in macrophages (Fig 3G, H). A recent study has reported that fibroblasts may inhibit the differentiation of oligodendrocyte precursor cells (46). Therefore, it is conceivable that reduction of the post-stroke fibrosis by Arg1 deletion in macrophages may influence the extent of the peri-infarct myelination. Indeed, we observed a decrease in the intensity of MBP immunoreactive signals in the peri-infarct region adjacent to the infarction border (Fig. 3I). The MBP density tended to be restored in the regions distant from the infarct boundary. Arg1 cKO animals showed significantly higher myelin density than the flox control group (Fig. 3I, J). The difference between the two groups was noticeable only in the regions within a couple of 100 μm from the border (Fig. 3I, K), suggesting that the fibrotic microenvironment may have a negative impact on the myelination of oligodendrocyte lineage cells that are positioned close to the fibrotic scar. Collectively, these findings suggest that Arg1 cKO in macrophages reduces post-stroke fibrosis accompanied by enhanced restoration of myelination in the peri-infarct cortex.

### Influence of Arg1 cKO in macrophages on synaptic structures and microglial synaptic elimination in the peri-infarct cortex

Synaptic structures undergo dynamic remodeling in the peri-infarct cortex (47, 48), and the synaptic plasticity in this region plays a crucial role in post-stroke functional recovery (49–52). Therefore, we reasoned that the notable improvement in motor functional recovery in Arg1 cKO animals may be linked to a differential regulation of synaptic connections in the peri-infarct region. The excitatory synaptic density was assessed by immunolabeling the excitatory pre-synaptic marker vGLUT2 and the post-synaptic marker PSD95 (Figure 4. A-E). Our analysis revealed a significant decrease in the number of both excitatory pre- and post-synaptic structures within the peri-infarct region at the 4-week time point, compared to animals that underwent a sham operation. The loss of pre- and post-synaptic markers was similar in extent. Importantly, the loss of excitatory synapses was prevented by depleting Arg1 in infiltrating macrophages (Figure 4B, F-H). When we accurately identified the synaptic contacts with overlapping signals between the pre- and post-synaptic markers using super-resolution imaging, the loss of synaptic contacts was more pronounced than either the pre- or post-synaptic marker alone (Fig. 4E, H). The reduction of the synaptic contacts was also alleviated by Arg1 cKO in macrophages. Together, these results suggest that Arg1-expressing macrophages infiltrating into the infarcted region may play a role in synaptic loss in the peri-infarct cortex, which could contribute to post-stroke functional deficits.

**Figure 4.**
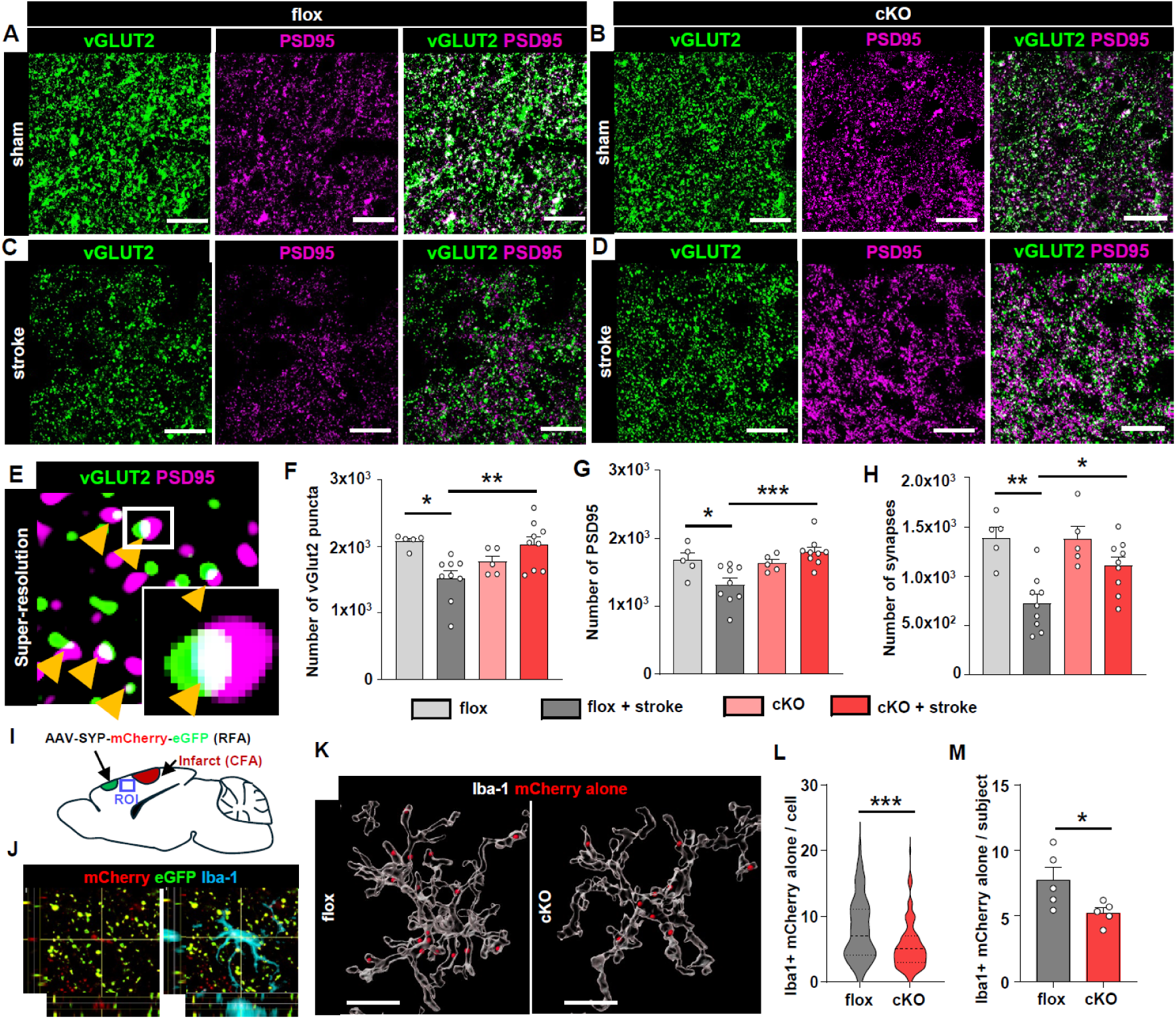
Influence of Arg1 cKO in macrophages on synaptic structures and microglial synaptic elimination in the peri-infarct cortex. (A-D) Representative images of excitatory pre-(vGLUT2) and post-synaptic (PSD95) markers in the peri-infarct region following a photothrombotic stroke. Scale bar = 5 μm. (E) Visualization of physical contacts between pre- and post-synaptic structures using super-resolution microscopic imaging. Colocalized (white) puncta were analyzed as synapses. (F-H) Quantification graphs comparing the number of pre-(F), post-synaptic puncta (G), and synaptic contacts (H). N = 28; flox + sham = 5, flox + stroke = 9, cKO + sham = 5, cKO + stroke = 9). *, **, and *** indicate p < 0.05, p < 0.01, and p < 0.001, respectively, by one-way ANOVA followed by post hoc Tukeys’s multiple comparisons. (I) An experimental scheme for analyzing *in vivo* microglial synaptic elimination using AAV9-Synaptophysin-mCherry-eGFP virus. Photothrombotic stroke was induced in the caudal forelimb area (CFA) 2 weeks after AAV9-SYP-mCherry-eGFP injection into the rostral forelimb area (RFA). Images were captured in the tissue adjacent to the infarction site to observe the dynamics of synaptic connections from the RFA to the peri-infarct region of CFA. (J) mCherry alone synaptic puncta detected in Iba-1 positive microglia. (K) 3D reconstructed Iba-1 and mCherry alone puncta. mCherry alone puncta inside the 3D rendered Iba-1 cellular membrane were converted to 3D spots using IMARIS software. Scale bar = 10 μm. (L) Violin plot illustrating the number of mCherry alone synapses per microglia. (N = 135 microglia from 5 animals for the flox group and 138 cells from 5 animals for the cKO group). *** indicates p < 0.001, by unpaired T-test. (M) The average number of mCherry alone synapses in Iba-1 positive cells. N = 5 for the flox group and 5 for the cKO group (16-32 cells per subject). * indicates *p* < 0.05 by unpaired T-test.

How does the deletion of Arg1 expressed in macrophages within the infarct affect synaptic structures in the peri-infarct cortex, located beyond the infarction border? The perineuronal nets (PNNs) are formed by fibrotic extracellular matrix (ECM) proteins containing several CSPG molecules and regulate synapse formation and neuronal plasticity (53, 54). PNNs undergo structural changes following a stroke, which may contribute to post-stroke functional recovery (55, 56). Therefore, it is possible that the post-stroke fibrosis, accompanied by an increase in CSPGs, could affect PNN remodeling. PNNs were visualized by Wisteria floribunda agglutinin (WFA), which binds to the PNNs surrounding Various interneurons. The number of PNN cells and the intensity of WFA signals were not changed in the peri-infarct cortex compared to sham-operated animals (Supplementary figure 5A-C). Furthermore, Arg1 cKO did not significantly influence the WFA signals in the peri-infarct cortex.

Recent studies suggest that microglial cells regulate the number of synapses in peri-infarct regions by phagocytosing synaptic structures (57, 58). We explored the possibility that microglial synaptic elimination in the peri-infarct cortex was affected by the deletion of Arg1 in macrophages within the infarcted tissue. To assess microglial synaptic elimination *in vivo*, we used AAV9-synaptophysin-mCherry-eGFP viruses to label neuronal synaptophysin. Given that the pKa of the red fluorescence proteins is lower than that of GFP, the mCherry signal is preserved in acidic environments such as in the phagocytic lysosome, whereas the GFP signal is rapidly dissipated (59). After delivery of AAV into the rostral forelimb area (RFA), we induced ischemic stroke in the caudal forelimb area (CFA) to examine synaptic connections originating from RFA to the adjacent ischemic region (Fig. 4I). mCherry alone signals observed within Iba-1 positive cells were determined as engulfed synapses by microglial cells (Fig. 4J). We quantified the number of the mCherry signals exclusively present in Iba-1 positive microglia using 3D rendered microglia and synapses (Fig. 4K). The violin plot showed that microglial cells from Arg1 cKO animals contained a significantly lower number of mCherry alone puncta than control floxed animals (Fig. 4L). The average number of mCherry alone puncta found within Iba-1 positive microglial cells was 7.7 ± 1.0 in control animals, and the average number was decreased to 5.2 ± 0.4 in Arg1 cKO group (Fig. 4M), revealing a significant reduction in microglial synaptic elimination by deletion of Arg1 in macrophages.

### Arg1 cKO in macrophages alters microglial inflammatory gene signatures in the peri-infarct cortex

To understand potential mechanisms underlying changes in microglial activity by Arg1 cKO in macrophages, we conducted cytokine PCR array to profile expressions of inflammation signature genes from FACS-sorted microglia (Fig. 5A). Microglia in the peri-infarct cortex was sorted based on CD11b immunoreactivity and low expression of monocyte marker CD45 (Supplementary fig. 6). A total of 53 cytokine genes were detected in microglial cells isolated from the peri-infarct cortical tissue, as listed in the heatmap plot (Figure 5B). The gene expression levels were compared between cKO and flox control animals in the stroke condition, and the fold change value was normalized by subtracting the difference between cKO and flox groups in the sham condition. All detected cytokine genes were plotted on a volcano plot to visualize the significant cytokine DEGs in Arg1 cKO compared to flox control animals following a photothrombotic stroke (Fig. 5C). Among the genes listed on the volcano plot, 10 were significantly downregulated, and only 4 genes were upregulated. Down DEGs included pro-inflammatory cytokines such as IL-6, IL-6 family oncostatin M (Osm), and TGF-beta with TGF-beta receptor ligands, including GDF9 and GDF15, while TGF-beta receptor antagonists such as BMP1 and BMP7 showed significant up-regulation. K-means clustering classified the significant DEGs into 4 clusters (Fig. 5D). Cluster 1 included proinflammatory cytokines such as IL-6, Osm, IL-1b, and IL-16. TGF-beta related genes, Tgfb1, Tgfbr1, GDF1, and GDF15 comprised of cluster 2. Cluster 3 consisted of BMP1, BMP7, and Fgf10, and TNF superfamily 12 and 13b (Tnfsf12 and Tnfsf13b), implicated in proinflammtory NFkB activation, were included in cluster 4. Interactions among the cytokines in each cluster were quantified using the STRING protein interaction analysis, which revealed IL-6 and TGF-beta as the top 2 cytokines with the highest interactions (Figure 5E). GO analysis showed that the cytokines DEGs were enriched in the pro-inflammatory pathway (regulation of interleukin-6 production, inflammatory response, interleukin-1-mediated signaling pathway) and the TGF-beta signaling (TGF-beta receptor signaling pathway, SMAD protein phosphorylation) (Figure 5F). Collectively, the gene expression study and bioinformatic analysis indicated that deleting Arg1 in macrophages infiltrating into the infarcted region reduced proinflammatory activity and TGF-beta signaling in microglial cells in the peri-infarct cortex.

**Figure 5.**
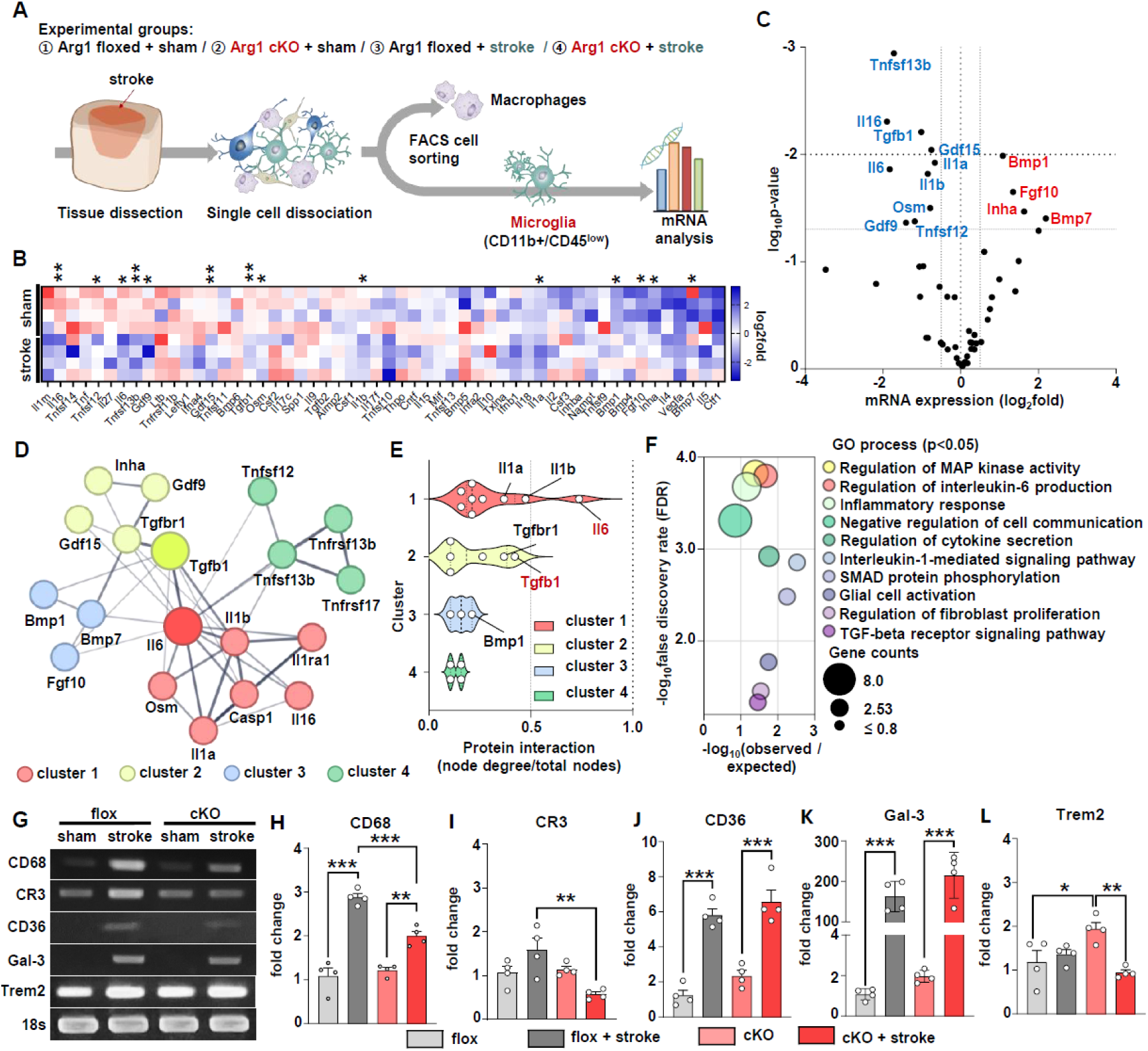
Arg1 cKO in macrophages alters microglial inflammatory gene signatures in the peri-infarct cortex. (A) A schematic diagram of the experimental design. A chunk of the cortical tissue, including the infarcted and spared peri-infarct cortex, was obtained, and the infarct core was carefully dissected and removed. The remaining peri-infarct cortical tissue was dissociated into single cells using the Adult brain dissociation kit, and the dissociated cells were subjected to FACS to isolate CD11b+/CD45^low^ microglial cells from macrophages. (B) A heatmap plot comparing all detected cytokine expressions between flox and cKO groups in the sham or stroke conditions. Log_2_(fold change) of the gene expression value of each cKO subject compared to the average expression value of the flox group was color-coded. Asterisks indicate the statistical significance of comparing cKO and flox groups in the stroke condition normalized by subtracting the difference between cKO and flox groups in the sham conditions. * and ** indicate *p* < 0.05 and *p* < 0.01, respectively, by unpaired T-test. (C) A volcano plot of DEGs by Arg1 cKO in the stroke condition. Significantly regulated genes were determined based on log_2_(fold change) > 0.5 and *p* < 0.05 criteria. Down DEGs were shown in blue, and up DEGs in red. The fold change value comparing cKO and flox control animals in the stroke condition was normalized by subtracting the difference between cKO and flox groups in the sham condition. (D) K-means clustering of the significant cytokine DEGs. (E) The degree of interactions within each cluster was calculated using the web-based STRING analysis. Protein interaction value was obtained by dividing the number of node degrees by the total number of nodes. (F) A bubble plot of the enriched GO processes (*p* < 0.05 = FDR < 95.0). The bubble size indicates the number of annotated genes in the GO process. (G) Representative images of semi-PCR products of microglial phagocytic markers (CD68, CR3, CD36, GAL-3, and TREM2). mRNA samples were obtained from FACS-isolated microglia out of the peri-infarct cortex 7 days post-stroke. (H-L) Quantitative comparison of mRNA expressions by real-time PCR. CD68 and CR3 showed significantly reduced expression in Arg1 cKO animals compared to flox animals after stroke. *, ** and*** indicate *p* < 0.05, *p* < 0.01 and *p* < 0.001, respectively, by One-way ANOVA followed by *post hoc* Tukey’s multiple comparison.

Since we observed attenuated synaptic eliminations by microglia in Arg1 cKO animals following stroke, we also examined the expression of phagocytosis-related genes Trem2, Gal3, CD36, CD68, and CR3 (60–64) in FACS-isolate microglial cells (Figure 5I). Among the phagocytic markers we measured, CD68, CD36, and Gal-3 were significantly up-regulated in microglia following stroke (Figure 5G-L). Arg1 cKO following stroke exhibited significantly down-regulated expression of CD68, but not CD36 and Gal-3 (Fig. 5H, J, K). CR3 expression was slightly increased in stroke conditions, and the level was significantly reduced by Arg1 cKO (Fig. 5I)

### Arg1 activation in macrophages influences microglia to enhance synaptosome phagocytosis *in vitro*

These findings suggest that macrophages infiltrating the infarcted tissue may interact with microglial cells in the peri-infarct cortex to regulate microglial activity. Recent studies also reported dynamic crosstalk between peripherally derived macrophages and microglia in CNS inflammation (19, 20). To directly test the interaction between the two cell types, we designed an *in vitro* experiment where bone marrow-derived macrophages (BMDMs) were co-cultured with microglial cells. To induce Arg1 activity in BMDMs, zymosan, a toll-like receptor 2 agonist, was treated based on a previous study (65), and zymosan treatment indeed markedly enhanced the Arg1 activity in BMDMs (Fig. 6A). The increased Arg1 activity was effectively abolished by Arg1 specific inhibitor OATD-02. Cultured BMDMs were treated with zymosan with or without OATD-2 for 24 hours and then co-cultured with microglia on a cell culture insert, allowing the two cell types to communicate with each other using diffusible molecules (Fig. 6B). Then, the cell culture insert with BMDMs were removed, and microglial cells were subjected to mRNA measurement or *in vitro* phagocytosis assay. We measured the expression of the two hub cytokine genes downregulated in the peri-infarct microglial cells following a photothrombotic stroke, TGF-beta1 and IL-6. Co-culturing microglia with zymosan-treated BMDMs did not alter the expression of TGF-beta1 (Fig. 6C). However, the mRNA levels of IL-6 were significantly upregulated in these co-cultures (Fig. 6D). The upregulation of IL-6 mRNA was attenuated when BMDMs in the co-culture system were treated with both zymosan and OATD-2 (Fig. 6D), indicating that the induction of IL-6 expression may be mediated by the enhanced Arg1 activity in BMDMs. To mimic microglial synaptic elimination, *in vitro* phagocytosis assay was performed using synaptosomes isolated from mouse cortical tissue. These synaptosomes were labeled with pHrodo, a pH-sensitive fluorescent dye that is expected to emit fluorescence when the synaptosomes are engulfed within an endosome at a low pH level (66). Microglial cells co-cultured with zymosan-treated BMDMs exhibited a higher number of pHrodo-positive cells compared to those co-cultured with PBS-treated BMDMs (Supplementary video and Fig. 6E), suggesting that BMDMs treated with zymosan enhanced the synaptic phagocytic activity of co-cultured microglial cells. When microglial cells were co-cultured with BMDMs treated with both zymosan and the Arg1 inhibitor OATD-02, the number of pHrodo-positive microglial cells decreased (Supplementary video and Fig. 6E). Quantitative analysis revealed that the zymosan + OATD-2 group showed a substantial reduction in pHrodo-positive cells during the early sessions of the assay, with a trend towards recovery in the later sessions (Fig. 6F). Quantification of the area under the curve (AUC) throughout the assay demonstrated a partial, yet statistically significant, reduction in the extent of synaptic phagocytosis by OATD-2 (Fig. 6G). These findings indicate that Arg1 activity in BMDMs influences synaptic phagocytosis observed in co-cultured microglial cells.

**Figure 6.**
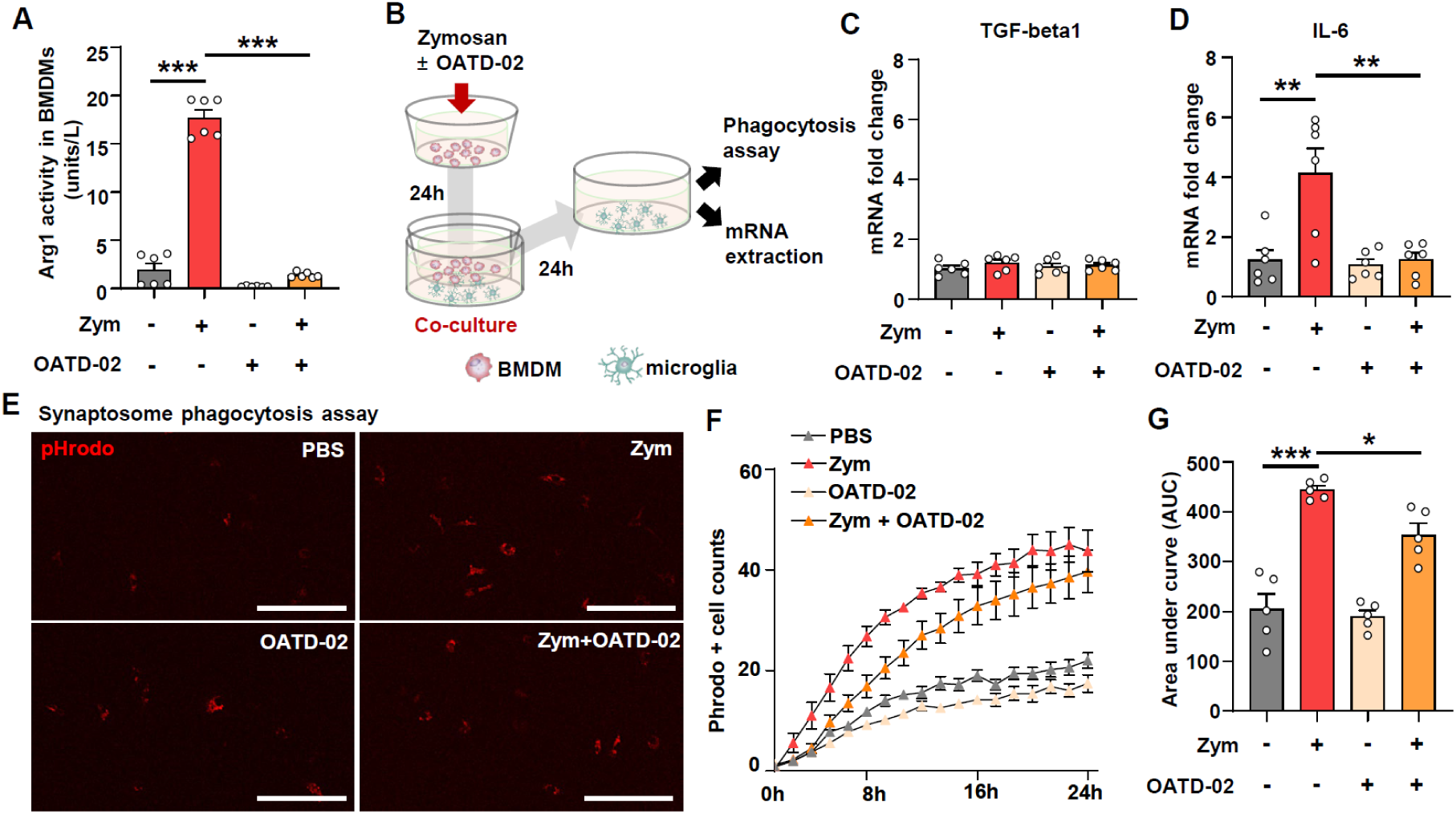
Arg1 activation in macrophages influences microglia to enhance synaptosome phagocytosis *in vitro*. (A) Arg1 activity assay in BMDMs following zymosan or OATD-02 treatment. *** indicates *p* < 0.001, by One-way ANOVA followed by *post hoc* Tukey’s multiple comparison. (B) A schematic diagram of the experimental design for the *in vitro* macrophage-microglia interaction assay. Bone-marrow derived macrophages (BMDMs) were cultured with Zymosan (1.25μg/ml) with or without Arg1 inhibitor, OATD-02 (1μM). After 24 hours, BMDMs were co-cultured with primary microglia for 24 hours. (C, D) Expression of TGF-beta1 (C) and IL-6 (D) mRNA expression in microglia at 24 hours after co-culture. ** indicates *p* < 0.01, by One-way ANOVA followed by *post hoc* Tukey’s multiple comparison. (E) Representative images of microglial pHrodo-synaptosome phagocytosis assay. Scale bar = 100 μm (F) Quantitative graph of pHrodo+ cell counts. (G) The area under cover of the graph (F) for statistical analysis. * and *** indicate *p* < 0.05 and *p* < 0.001, respectively, by One-way ANOVA followed by *post hoc* Tukey’s multiple comparison.

## Discussion

The current study investigated the functional role of Arg1, a well-established marker for anti-inflammatory macrophage phenotype, in the context of ischemic stroke. Deletion of Arg1 in LysM-positive macrophages resulted in substantial improvement of skilled forelimb motor functions, suggesting a detrimental influence of macrophage-expressed Arg1 on post-stroke functional recovery. Arg1 cKO in macrophages attenuated fibrotic scar formation accompanied by restoration of myelination in the peri-infarct cortex. Furthermore, animals with Arg1 cKO showed an increase in synaptic density and diminished microglial synaptic elimination in the peri-infarct region, an area distant from the infarction core. Gene expression analysis revealed that Arg1 deletion in macrophages led to reduced TGF-β signaling and pro-inflammatory cytokine activity. An *in vitro* co-culture experiment with macrophages and microglial cells demonstrated dynamic interactions between these cell types, wherein arginase activity in macrophages regulated microglial synaptic phagocytosis.

Arg1 expression has been documented in infiltrating macrophages or monocyte-derived macrophages in various CNS injuries, including ischemic stroke, spinal cord injury, and demyelinating diseases (28, 67, 68). Traditionally, Arg1 expression has been associated with improved functional outcomes (28, 69) and is often considered a marker for macrophages that promote CNS tissue repair or regeneration (68, 70, 71). However, few studies have directly addressed the functional role of Arg1 expression in macrophages within these contexts. A recent study demonstrated that depletion of Arg1-positive microglia/macrophages using mannosylated clodronate liposomes exacerbated post-stroke outcomes by promoting proinflammatory responses (72). However, this study examined the functional roles of Arg1-positive myeloid cells, rather than the Arg1 protein itself. Another study reported that knockout of STAT6 led to increased infarction volume and worsened functional outcomes, partly by suppressing Arg1 expression in macrophages and microglial cells. Lentiviral overexpression of Arg1 restored the anti-inflammatory features (73), supporting a beneficial influence of Arg1 on post-stroke recovery. To our knowledge, the present study is the first to directly examine the functional role of Arg1 in macrophages in an animal model of ischemic stroke. Contrary to the prevailing idea that Arg1-positive macrophages contribute to the resolution of inflammation and/or tissue regeneration following CNS injuries, our findings suggest a potentially detrimental role for macrophage-specific Arg1 in post-stroke recovery, challenging current paradigms of Arg1 function in CNS injury.

Our cellular mapping study has identified LysM-positive infiltrating macrophages as a primary source of Arg1 following photothrombotic stroke. This finding was corroborated by a marked reduction in Arg1-positive cells upon Arg1 deletion using the LysM-Cre line. These results align with previous research demonstrating exclusive Arg1 expression in LysM-GFP-positive cells, but not in GFP-negative myeloid cells, in the middle cerebral artery occlusion model (74). We observed that approximately 20% of Arg1-positive cells colocalized with CX3CR1-GFP-positive cells. These cells may represent either genuine resident microglia or microglia-like cells derived from infiltrating hematogenous macrophages. In the latter case, Arg1 in these microglia-like cells would be deleted by the LysM-Cre line, given their hematogenous macrophage origin (32). It is noteworthy that not all Arg1-positive cells were LysM-positive, and a small fraction of Arg1-expressing cells persisted in Arg1 cKO animals. This residual population may represent a very minor subset of resident microglial cells expressing Arg1. Alternatively, it could indicate that not all infiltrating macrophages are LysM-positive, as previous studies have reported that a small population of infiltrating macrophages can express CX3CR1 (75). Regardless of their origin, we speculate that the influence of the small amount of residual Arg1 is likely negligible in the context of our study.

Our study demonstrates that deletion of Arg1 in infiltrating macrophages substantially reduced the accumulation of fibrotic extracellular matrix (ECM) both in the infarcted tissue and at the border between the infarction core and peri-infarct cortex. While fibrosis is a key component of wound healing, excessive fibrosis can impede functional restoration following injury in various organs. In CNS, fibrotic scarring has been shown to inhibit axonal regeneration and diminish functional recovery (44, 76). In stroke models, reduction of fibrotic scarring through pharmacological interventions has been associated with increased peri-infarct axonal sprouting and improved functional recovery (37, 38). In our study, Arg1 deletion did not affect the sprouting of 5-HT axons or the extent of axonal injury in the peri-infarct cortex. Instead, Arg1 cKO enhanced myelination in the peri-infarct cortex close to the infarction border. A recent study revealed that photothrombotic stroke led to demyelination in the peri-infarct cortex, which could be improved by regulating inflammatory signaling (77). We also observed decreased myelination in areas of excessive fibrotic ECM accumulation near the infarction. This pattern is reminiscent of the experimental autoimmune encephalomyelitis (EAE) model, where perivascular fibroblast infiltration in the spinal cord parenchyma correlates with areas of demyelination (46). These findings suggest that fibrotic ECM accumulation, orchestrated by Arg1 in infiltrating macrophages, may inhibit remyelination by oligodendrocyte lineage cells in the peri-infarct cortex close to the infarction border. Arg1 in macrophages promotes fibrotic scar formation in cutaneous injuries by facilitating collagen synthesis through proline production (26). In our study, we found that deleting Arg1 in LysM-positive macrophages reduced TGF-beta signaling, a crucial pathway regulating fibrosis, in microglial cells in the peri-infarct cortex. This decrease in TGF-β activity in microglia near the infarction border may also contribute to the reduced fibrotic scarring observed in Arg1 cKO mice.

Immunolabeling of excitatory synapses revealed that photothrombotic stroke significantly reduced the number of excitatory synapses in the peri-infarct area. Notably, depletion of Arg1 in infiltrating macrophages mitigated this synaptic loss. Microglia are known to play a crucial role in regulating synapse formation in the brain through synaptic pruning during developmental stages (78). In neurodegenerative diseases like Alzheimer’s disease, microglial immune responses lead to the elimination of viable synapses, exacerbating synaptic loss (79). Similarly, synaptic elimination by microglial phagocytosis also occurs in the peri-infarct region following stroke (57, 58). In these studies, inhibition of synaptic phagocytosis improved post-stroke functional recovery, suggesting that microglia-mediated loss of synaptic structures may contribute to neurological deficits following stroke. Therefore, it is highly likely that decreased microglial synaptic elimination by Arg1 cKO in the current study contributed to the improvement of motor recovery. We found reduced proinflammatory cytokine activity and TGF-beta signaling in microglial cells from Arg1 cKO animals, suggesting that pro-inflammatory cytokines and TGF-beta signaling in the peri-infarct cortex may drive microglial synaptic phagocytosis, resulting in decreased synaptic density. Microglial states regulating synaptic phagocytosis are influenced by various inflammatory cytokines (80). Recent research reported that SPP1-mediated microglial synaptic elimination is regulated by autonomous TGF-beta signaling in an Alzheimer’s disease model (81), implicating TGF-beta signaling in the activation of microglial synaptic elimination.

Our *in vitro* experiment directly addressed the question of how Arg1 deletion in infiltrating macrophages in the infarcted tissue alters the post-stroke inflammatory milieu in the peri-infarct cortex. We observed that zymosan-treated BMDMs significantly enhanced synaptic phagocytosis in co-cultured microglial cells. Notably this enhanced phagocytic activity was attenuated by co-treatment with the Arg1 inhibitor OATD-02, suggesting that increased Arg1 activity in BMDMs was responsible for the heightened phagocytic response in microglia. These findings imply that macrophages infiltrating the infarcted tissue exert modulatory influence on microglial cells in the peri-infarct tissue. The precise mechanism by which macrophages modulate microglial phagocytic activity warrants further investigation. Modulation of microglial phagocytosis by peripherally derived macrophages was also observed in a spinal cord injury model (19). Conversely, microglial cells can regulate the recruitment and neurotoxic effects of monocyte-derived macrophages (21). These observations suggest a bidirectional interaction between macrophages and microglial cells, which may play a crucial role in shaping the post-injury inflammatory milieu (20).

In conclusion, our study elucidates the detrimental role of Arg1 expressed in macrophages following ischemic stroke. We demonstrate that Arg1 deletion in infiltrating macrophages confers beneficial effects on post-stroke recovery through two primary mechanisms: regulation of fibrotic scar formation and modulation of microglial synaptic phagocytosis. These findings suggest that Arg1 in macrophages represents a promising therapeutic target for modulating the post-stroke inflammatory environment. However, given the constitutive expression of Arg1 in the liver, systemic inhibition of Arg1 could potentially lead to hepatotoxicity. To circumvent this issue, we propose that infusion of macrophages preconditioned with *ex vivo* Arg1 inhibition could be exploited as a targeted approach to mitigate Arg1 activity in infiltrating macrophages following stroke.

## Acknowledgements

This research was supported by the National Research Foundation of Korea (NRF) research programs (2021R1A2C2006110, RS-2023-00244748, 2019R1A5A2026045 to BGK), and the NRF Global Ph.D. Fellowship Program (2018H1A2A1061966 to HSK). The Arg1 inhibitor OATD-02 was produced by Molecure S.A. in Poland under a material transfer agreement (MTA).

## Author Contributions

The first author Hyung Soon Kim conducting the majority of the research.

