## Supplementary Figures for "Detrimental Influence of Arginase-1 in Infiltrating Macrophages on Post-Stroke Functional Recovery and Inflammatory Milieu"

### Supplementary Data

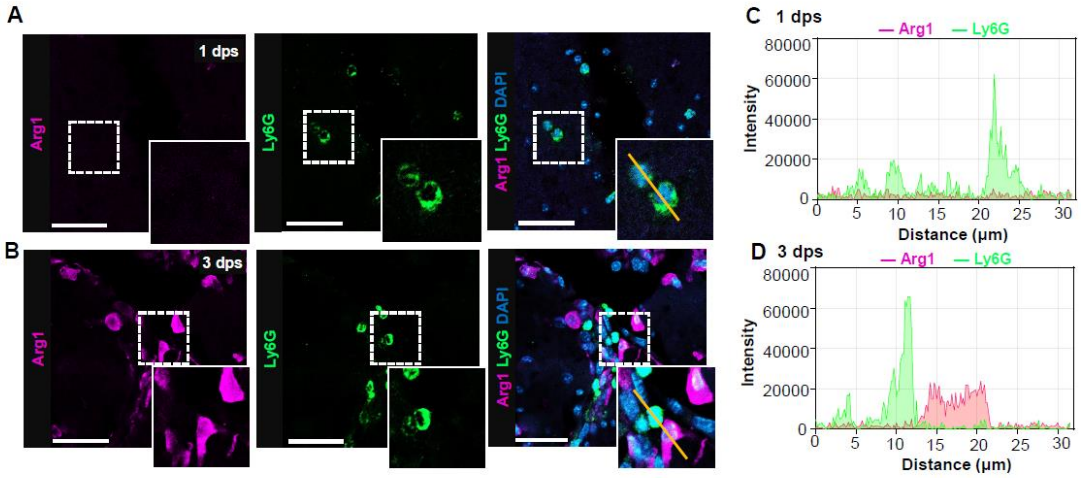


### Supplementary figure 1.

(A, B) Representative images of the immunohistochemical staining of Arg1 and Ly6G at 1- (A) and 3-days (B) post-stroke (dps). The white dotted squares indicate the magnified regions of the inset images. Scale bar = 50 μm. (C-D) Histogram of fluorescence intensity of the yellow solid lines in the representative images at 1 (C) and 3 (D) dps, respectively.

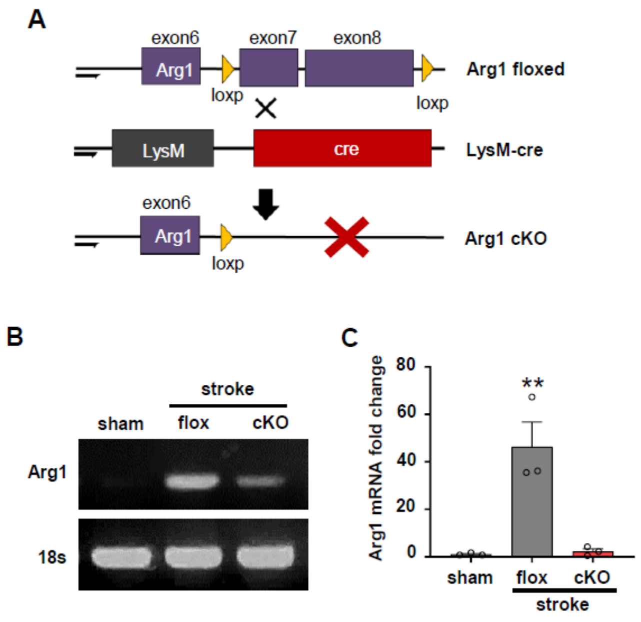


**Supplementary figure 2.**

(A) A schematic depiction of the Cre-loxP conditional knockout system. The loxP sequences (yellow arrowheads) are inserted downstream of exon 7 and upstream of exon 8 of the Arg1 genomic sequences. LysM-cre animals express Cre-recombinase under the control of the Lysozyme2 (LysM) promoter so that Arg1 is deleted only in cells expressing LysM. (B) Electrophoresis of PCR-amplified Arg1 from the cortical tissue from sham-operated animals or animals after a photothrombotic stroke with either flox and cKO genotype. (C) Real-time PCR result of Arg1 mRNA expression in the cortical tissue. ** indicates $p < 0.05$ by two-way ANOVA followed by *post hoc* Bonferroni's multiple comparisons. N = 3 animals per group.

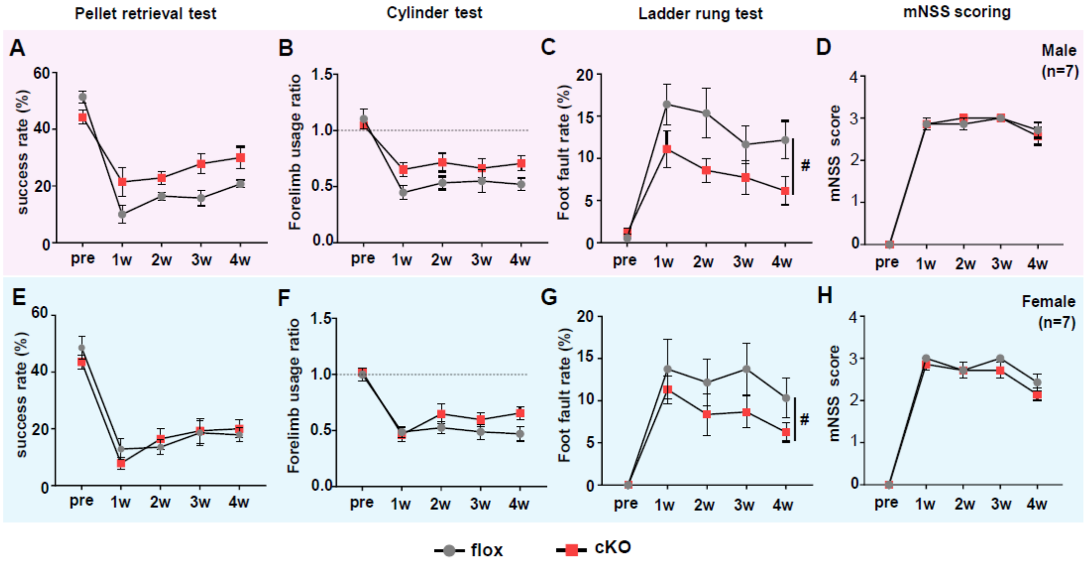


**Supplementary figure 3.**

(A-D) Behavioral analysis of each task only in males, or (E-H) females. Two-way ANOVA repeated measure was performed to match the time difference. Bonferroni's multiple comparison was conducted to assess group differences at each time point. Both males and females showed similar significance in the cylinder and ladder walk tests, indicating no sex difference in functional recovery after stroke in Arg1 cKO animals. (N = 7 per sex). # indicates $p < 0.05$ by two-way ANOVA column factor (overall group difference).

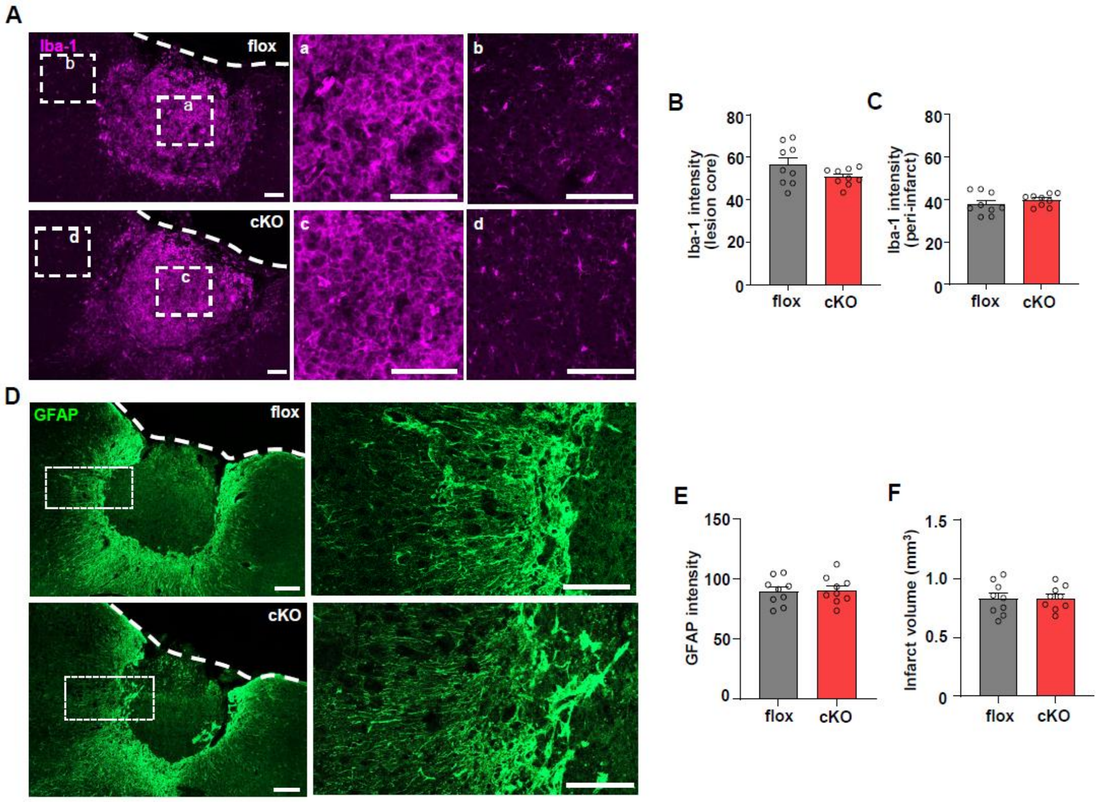


**Supplementary figure 4.**

(A) Representative images of Iba-1 positive cells in the infarcted cortex area 4 weeks after a photothrombotic stroke. Dashed rectangles indicate regions that are magnified on the right side. Dashed lines indicate the outer edges of the cerebral cortex. Scale bars = 50 µm. (B, C) Quantification graphs comparing Iba-1 immunoreactivity at the lesion core (B) and peri-infarct cortex (C). (D) Representative images of GFAP immunostaining in the infarcted cortex area 4 weeks after a photothrombotic stroke. . Dashed rectangles indicate regions that are magnified on the right side. Dashed lines indicate the outer edges of the cerebral cortex. Scale bars = 100 µm. (E, F) Quantification graphs comparing the intensity of GFAP immunoreactivity (E) and the volume of infarcted tissue delineated by GFAP positive astroglial border (F).

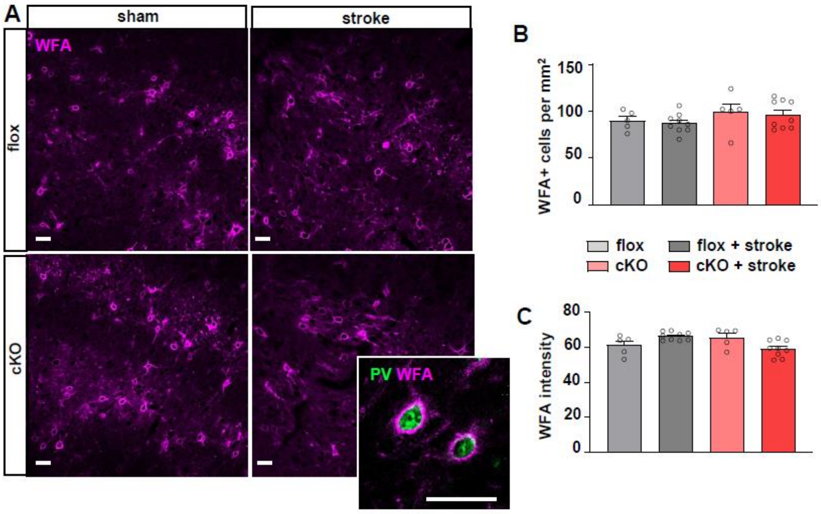


**Supplementary figure 5.**

(A) Wisteria floribunda agglutinin (WFA) labeled peri-neuronal nets (PNNs) in the peri-infarct region of Arg1 flox control and cKO animals with stroke, or in the anatomically same region of the cortex of sham-operated animals from flox control and cKO animals. The inset image shows parvalbumin (PV) expressing interneurons surrounded by WFA positive PNNs. Scale bar = 50 μm. (B, C) Quantification graphs comparing the number of WFA positive neurons (B) and the intensity of WFA signals (C) in the peri-infarct cortex.

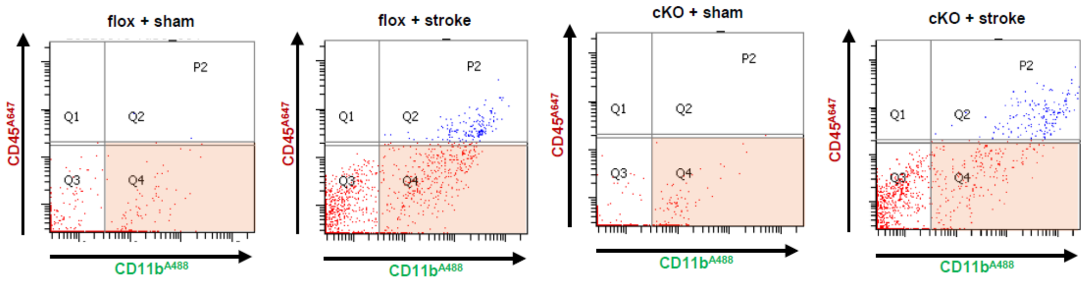


**Supplementary figure 6.**

Fluorescence-activated cell sorting (FACS) gating strategies to isolate microglia from the peri-infarct cortex. After single-cell dissociation using an adult brain dissociation kit, cells were labeled with CD45-Alexa647 and CD11b-Alexa488 antibodies. $CD45^{low}CD11b^{+}$ Q4 fractions (in red) were sorted as microglia fractions for all groups. Isolated microglia fractions were used for further analysis.

**Supplementary video 1.**

Live images of microglial pHrodo-synaptosome phagocytosis assay. Each frame of the video represents 80 minutes, and live imaging was conducted for 24 hours immediately following pHrodo-synaptosome treatment. Microglia were incubated for 24 hours with conditioned media collected from a macrophage and microglia co-culture before pHrodo-synaptosome treatment. Scale bar = 50 μm.
